# PPI-ID: Streamlining Protein-Protein Interaction Prediction through Domain and SLiM Mapping

**DOI:** 10.1101/2025.04.22.649917

**Authors:** Haley V. Goodwin, Nigel S. Atkinson

## Abstract

AlphaFold-Multimer models protein complexes and facilitates protein-protein interaction (PPI) prediction. Mapping of protein interaction domains and motifs onto the 3D structure can lend credence to the model and provide insight into the function of a given interaction. Furthermore, limiting structure prediction to only the domains and motifs that are likely to interact can reduce the computational demand and produce a higher quality model. To satisfy these needs, we built the Protein-Protein Interaction Identifier (PPI-ID). PPI-ID maps interaction domains and motifs onto molecular structures and filters for those that are sufficiently close to interact. Once an interface is found, PPI-ID labels interacting amino acids. Given only sequences, PPI-ID predicts regions for AlphaFold-Multimer modeling, reporting potential interactions only when each protein has one-half of a paired sequence. Testing with known dimers confirms high accuracy of the tool.

## INTRODUCTION

Many protein-protein interaction (PPI) domains and interaction motifs have been archived in databases. Domain-domain interactions (DDIs) between large structured regions are important for stable protein multimerization, while short linear motifs (SLiMs), often located in intrinsically disordered regions (IDRs), are thought to mediate short-lived signaling interactions and to play a lesser role in stable protein interactions (1).

Systematic curation and annotation of known protein domains and motifs have enhanced our ability to understand protein interactions, signaling pathways, regulatory pathways, and disease mechanisms (2, 3). The shortest of these are SLiMs, which are 3-10 amino acids long and defined by regular expressions. SLiMs are difficult to physically study because of apparent weak affinity for and transient interaction with their targets (1). In addition, prediction algorithms overpredict these sequences in proteins, reducing their utility to indicate protein function. When interpreting the meaning of a SLiM in a protein, it is best to have supporting evidence that the motif is important. This usually involves physical protein-protein interaction assays, evolutionary conservation analysis, mutation analysis, and, very recently, mapping of the motif onto models of interacting proteins (4).

AlphaFold-Multimer can be used to predict protein structure and interactions (5, 6). Mapping interaction domains and motifs onto an existing PPI model (top-down approach) can lend credence to the model or provide insight to the function of a given interaction. Analyzing primary protein sequences for domains and motifs can also be useful for selecting which protein regions should be modelled (bottom-up approach). Bret et al. and Lee et al. showed that multimeric predictions can be improved if one focuses on sequences involved in the interface formation, thereby decreasing confounding intramolecular contacts (7, 8).

Mapping interaction domains and SLiMs onto proteins is conveniently achieved using the InterPro and ELM websites (https://www.ebi.ac.uk/interpro/ and http://elm.eu.org/, respectively) (2, 9). However, neither nor ELM can confirm that two proteins contain an appropriately matched complement of domains and/or motifs in the appropriate positions to interact. Manually evaluating these criteria can become time consuming and subject to error, especially if a protein has a large number of domains and/or motifs or if many protein pairs need to be surveyed. For the top-down approach, it would be helpful if matched pairs could be displayed on a 3D molecular model and if potential interfaces could be filtered for domains and motifs close enough to interact. For the bottom-up approach, it would be ideal if a program reported matches only if each protein contained one-half of a pair of interaction sequences. To facilitate these approaches, we generated the Protein-Protein Interaction Identifier (PPI-ID). The program can be accessed at http://ppi-id.biosci.utexas.edu:7215/.

## METHODS

### Required Libraries

PPIP-ID is written in R with a web interface generated using Shiny. The InterPro, ELM, and UniProt database APIs, were accessed with *httr*. The *r3dmol* and *bio3d* packages were used to import, format, and display pdb information (10, 11).

### Database Composition

The PPI-ID database of 34,189 unique domain-domain interactions (DDIs) was compiled from the 3did (2022 release) and DOMINE (2010 release) databases (https://3did.irbbarcelona.org/index.php and https://manticore.niehs.nih.gov/cgi-bin/Domine, respectively) (3, 12). 3did provides DDIs mapped in crystal structures. DOMINE provides a combination of crystal structure and high-confidence predicted DDIs. A database of 399 DMIs based on crystal structures were gathered from the ELM Database (2024 release) (2). Domain information is stored according to Pfam ID. Motif information is stored according to ELM classifiers.

### Extracting Domain/Motif Information

A number of functions query the compiled DDI/DMI databases and predict potential interactions between a pair of user-provided proteins. These functions then check Pfam or ELM IDs against DDI/DMI databases to determine whether a pair of domains or a domain and a motif constitute a potential interaction. If so, the pair, along with the associated amino acid ranges for the domains or motif are documented.

#### DDI Prediction

Within the ‘Predict from Accession’ tab, accession numbers are used to access the InterPro API (
https://www.ebi.ac.uk/interpro/api/entry/pfam/protein/uniprot/[Accession Number]. Within the ‘Predict from Sequence’ tab, user-provided TSV files obtained from InterProScan are analyzed.

#### DMI Prediction

Within the ‘Predict from Accession’ tab, accession numbers are used to access the InterPro or the UniProt API for domain and fasta information, respectively. The UniProt API URL used within PPI-ID to fetch appropriate amino acid sequences is https://rest.uniprot.org/uniprotkb/[Accession Number]. The protein sequence is searched for the presence of a SLiM using the str_locate_all() function from the *stringr* package.

Within the ‘Predict from Sequence’ tab, user-provided TSV files from InterProScan and ELM Predict are analyzed. The user can also obtain SLiM information from PPI-ID directly, using the ‘Get SLiM Information’ tab. PPI-ID is capable of predicting DDIs or DMIs, but not motif-motif interactions, because available databases do not document motif-motif interactions. If a user attempts to seek motif information for both protein inputs, PPI-ID will return an error and prompt for domain information for at least one protein.

### Filter by Contact Distance

If the user has provided a pdb file of the protein complex, the table of predicted DDIs/DMIs can be filtered for contact distance (in angstroms). Contact distance filtering is done by the filter_by_distance() function, which uses the atom.selection() and cmap() functions from the *bio3d* library to select Cα’s and determine whether DDIs/DMIs are within the user-provided contact distance.

### PPI-ID Validation

A total of 80 PPI complexes were folded using AlphaFold-Multimer (version 2.3.2), with 40 folds for DDI validation and 40 folds for DMI validation. AlphaFold-Multimer was run at the Texas Advanced Computing Center (TACC) using the reduced database. A total of 5 machine learning models were produced with a single pdb prediction per model.

For DDI validation, 40 PDB database entries were randomly selected from a collection of 14,972 crystal structures containing DDIs, as curated by 3did (3). Only PDB database entries representing inter-protein interactions were selected. Entries containing synthetic proteins were excluded. For bottom-up validation, the accession numbers were input into PPI-ID. For top-down validation, contact distance filters of 5 to 7 angstroms were used. In both cases, program output was checked to confirm that the interacting domains were correctly identified. For DMI validation, 40 PDB database entries were randomly selected from a collection of 923 crystal structures containing DMIs, as curated by Bret et al. (7). The previously described bottom-up and top-down procedures used for DDI validation were used. Validation data can be found in Supplementary Table 1 and Supplementary Table 2, respectively.

## RESULTS

PPI-ID facilitates structural analyses of predicted dimeric models through molecular visualization, contact distance-based interface identification, and contact residue labelling. PPI-ID predicts potential interaction interfaces from protein sequence information using information from ELM, 3did, Interpro, and DOMINE databases (2, 3, 9, 12). In the top-down approach (Figure 1, right), PPI-ID graphically maps the Domain:Domain Interactions (DMI) and Domain:Motif Interactions (DMI) onto the 3D model of the protein dimer only if each protein contains compatible interaction sequences and only if these sequences are within a specified distance of one another. For the bottom-up approach (Figure 1, left), PPI-ID displays the position of DDI and DMI sequences in the primary amino acid sequence for two proteins only when each protein contains one-half of a pair of protein:protein interaction signature sequences. It expresses these as a table of the amino acid residue numbers from each protein that are predicted to interact (Figure 2). This information might be used to select the amino acid residues to be used to model the interacting regions of the proteins.

**Figure 1.**
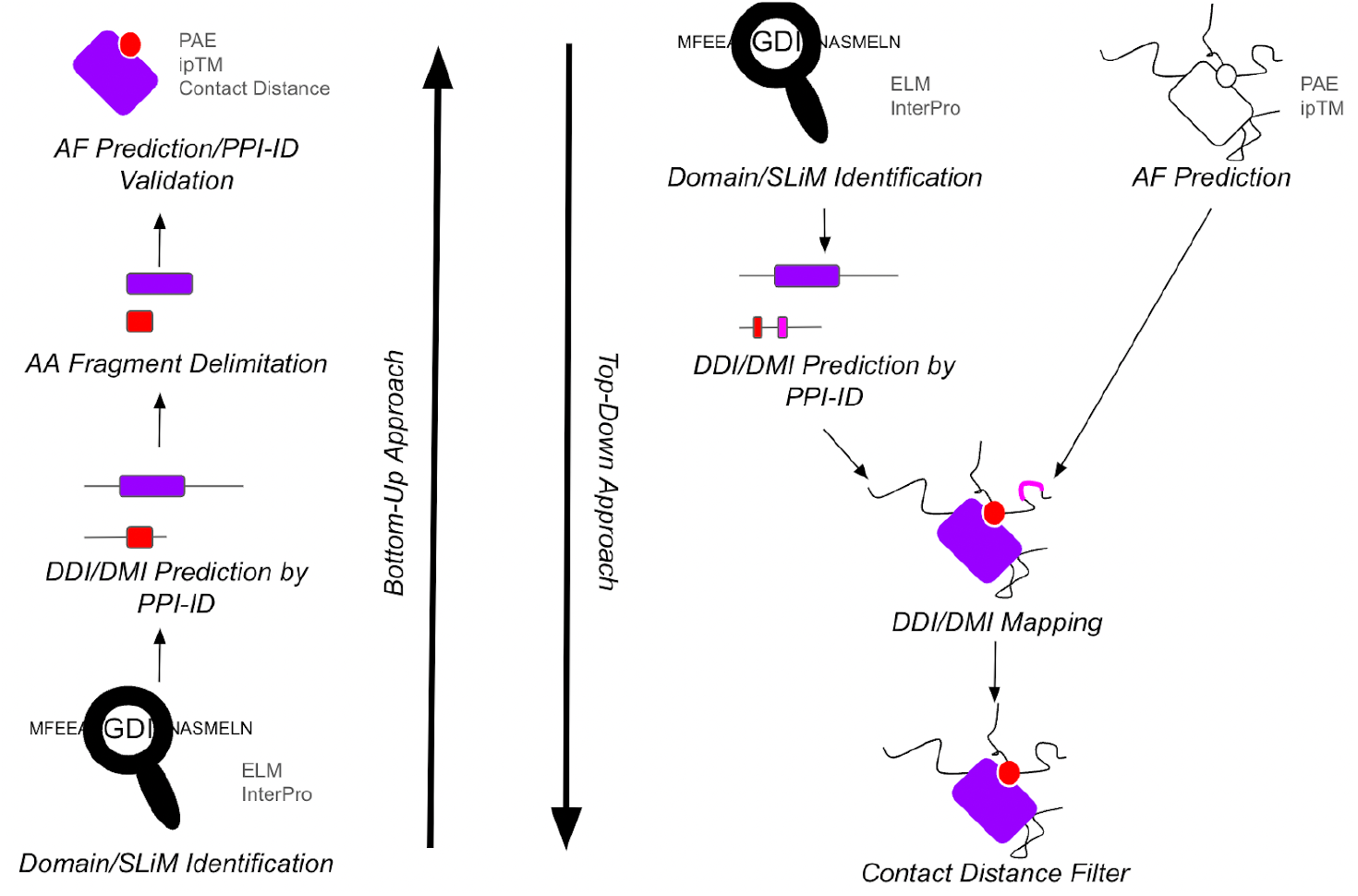
Summary of analyses by PPI-ID. ***Left side:*** The bottom-up approach searches two protein fasta files for domains and SLiMs in each protein that could facilitate a physical interaction between the two proteins. Output is presented in a way that facilitates cropping of the fasta files before using AlphaFold-Multimer which can favor the generation of a higher accuracy model of the region. ***Right side:*** In the top down approach, protein-protein interaction domains/SLiMs are mapped onto a 3D heterodimer model of two proteins and their positions visualized. Domains or SLiMs that are too far apart to interact can be filtered out thereby enriching for sequence pairs that have the greatest potential for a physical interaction.

**Figure 2.**
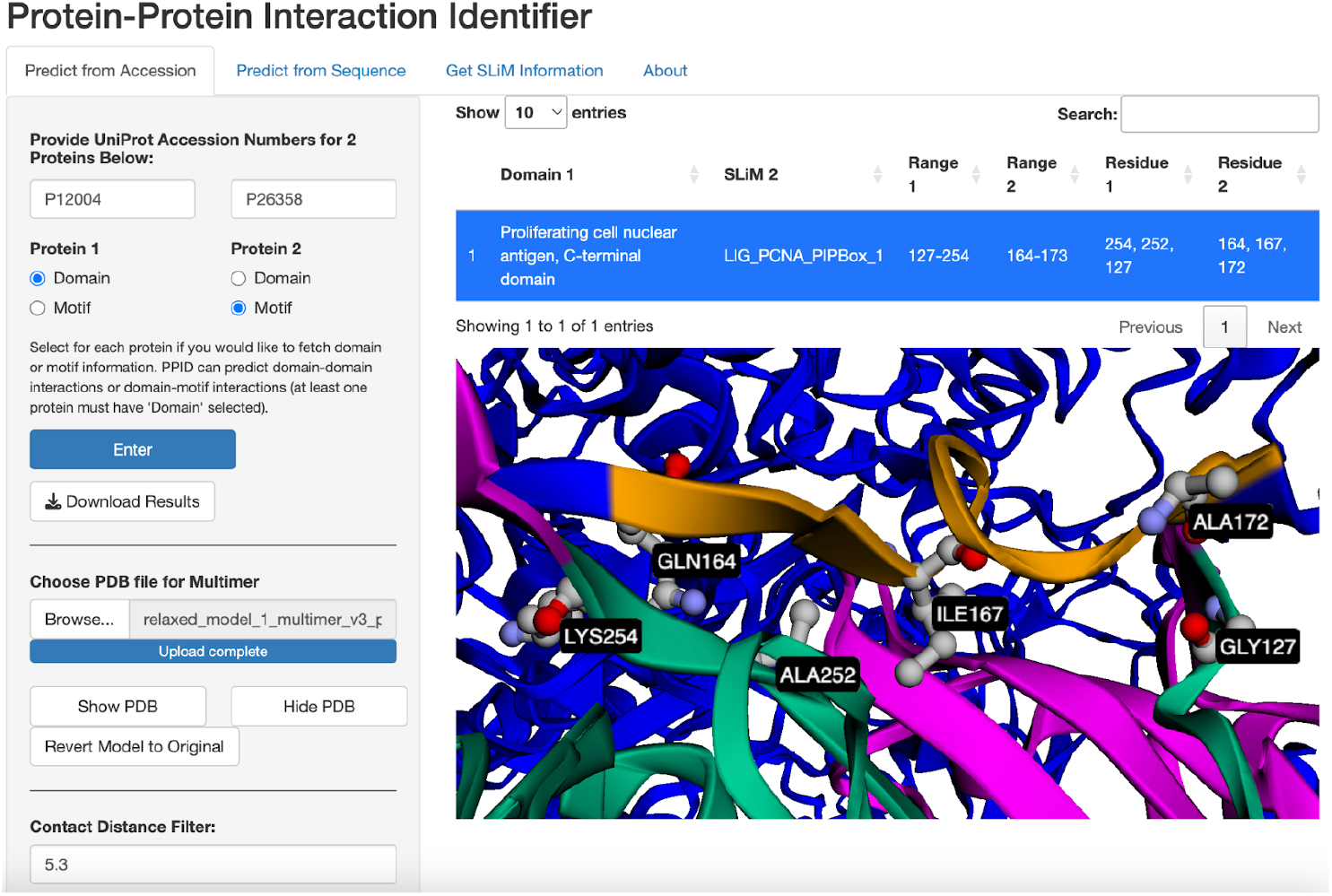
The top-down approach using the human PCNA (P12004) and DNMT1 (P26358) proteins. *Contact Distance Filter* was used to only show predicted interactions that come within 5.3 Å. The region selected in the table is displayed on the protein in green and yellow (PCNA and DNMT1, respectively). The *Add Contact Labels* button (not pictured) labels residues within the distance filter limit and represents them in ball-and stick form with carbon in gray, oxygen in red, and nitrogen in light blue.

Case studies presented in this paper take advantage of the ‘Predict from Accession’ tab within PPI-ID. However, in the ‘Predict from Sequence’ tab, users are also able to upload domain information for a protein (obtained from InterProScan) and motif information for a protein (obtained using the ‘Get SLiM Information’ tab of PPI-ID). The benefit of uploading annotated sequence information in the ‘Predict from Sequence’ tab is that user predictions are not limited to proteins with associated UniProt accession numbers. As a result, users can perform computational PPI prediction and analysis with synthetic proteins, uncharacterized isoforms, and protein fragments.

### PPI Prediction

#### PCNA and DNMT1 Interaction, Top-Down Example

For the top-down approach, PPI-ID presents a detected DMI between human proliferating cell nuclear antigen (PCNA) and DNA methyltransferase 1 (DNMT1) proteins (Figure 2). Many PCNA pairing partners contain a conserved motif called the PCNA-interacting protein (PIP) box (13–15). One PIP box-containing protein that has been demonstrated to interact with PCNA is DNMT1 (16). InterProScan sequence alignment shows the PCNA amino acid sequence contains 20 domain entries. DDI and DMI are stored by Pfam ID or ELM ID, so PPI-ID only analyzes entries using these identifiers. A search of the PCNA sequence identifies two Pfam domains: the PCNA C-terminal domain and the PCNA N-terminal domain. Regular expression searches on the DNMT1 amino acid sequence identifies 546 possible SLiMs. Manually assessing the compatibility of every possible domain:motif pairing would require the manual inspection of 1092 pairs.

PPI-ID automates the identification of potential interactions between the PCNA C-terminal domain and two PIP box motifs in DNMT1. When a 5.3 Å contact distance filter and contact residue labelling are applied, PPI-ID identifies and highlights residues likely important for mediating interactions between PCNA and DNMT1 – specifically Ala252 and Gly127 of PCNA which are known to hydrogen bond with Gln164 and Ala172 of DNMT1, respectively (16). Furthermore, Ile167 of DNMT1, also identified by the contact filter, has been shown to plug into the hydrophobic binding pocket of PCNA (16).

#### DifA and Cactus Interaction, Bottom-Up Example

For the bottom-up approach, PPI-ID displays the position of signature sequences for DDIs or DMIs that could contribute to an association between the two proteins. The *Drosophila* DifA and Cactus proteins are well-known interacting partners, with this interaction taking place between the Rel homology region dimerization domain of DifA and the ankyrin repeats of Cactus (17). Figure 3 shows the result of submitting the accession numbers of DifA and Cactus proteins for DDI prediction. Namely, the DNA-binding and dimerization domains of the DifA Rel-homology region are identified as potential interactors with ankyrin repeat domains found within Cactus. This information can be used to delimit the protein sequences for AlphaFold-Multimer modeling, which enhances the accuracy of the structure (7, 8).

**Figure 3.**
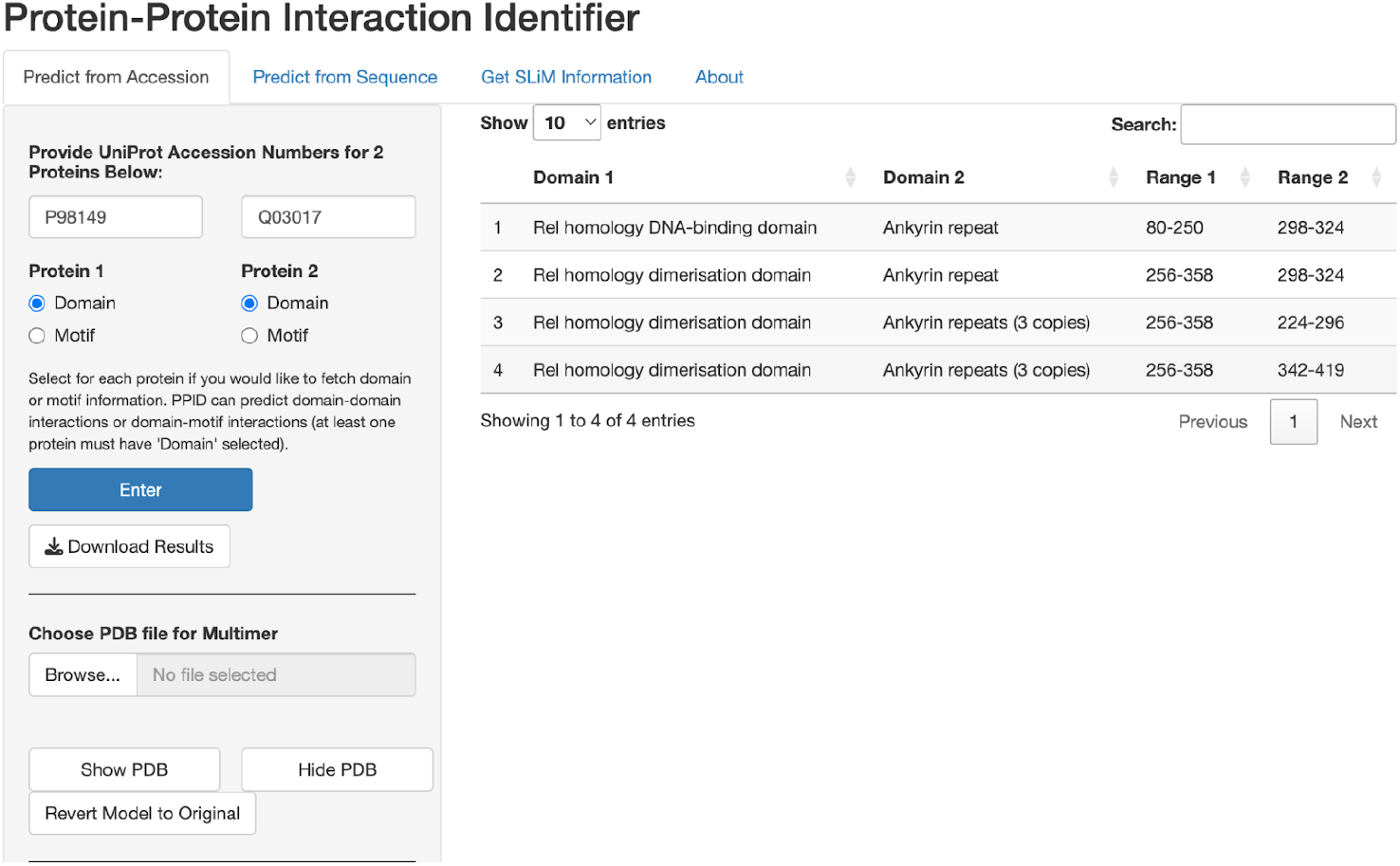
The bottom-up approach using the DifA (P98149) and Cactus (Q03017) proteins, which are known to bind. The table shows four predicted interactions that take place between ankyrin repeats of Cactus and the Rel homology domain of DifA. Relative positions of each domain are shown.

### PPI-ID Validation

80 PPIs with documented crystal structures were folded with AlphaFold-Multimer. 40 PPIs were used to validate DDI prediction accuracy, and the remaining 40 were used to validate DMI prediction accuracy. The accuracy of PPI-ID top-down and bottom-up analyses is summarized in Table 1.

**Table 1.** Percentage of known DDIs and DMIs successfully identified. Forty protein pairs were used to validate DDI identification capability, and 40 protein pairs were used to validate DMI identification capability.

| Interaction Type | Bottom-Up % Correct | Top-Down % Correct |
| --- | --- | --- |
| Domain-Domain (DDIs) | 98% | 85% |
| Domain-Motif (DMIs) | 95% | 73% |

#### Top-Down Validation

Pairs of protein accession numbers and pdb files of predicted complexes were submitted to PPI-ID. A prediction was considered successful if the proper interaction interface was identified both in PPI-ID output and in the AlphaFold-produced structure. PPI-ID top-down validation yielded respective accuracy rates of 85% and 73% for DDIs and DMIs. Using a similar top-down approach, Bret et al. reported that AlphaFold-Multimer had a predictive performance of 52.7% (7). The increased success of PPI-ID over AlphaFold-Multimer analysis alone is likely a product of sampling error.

#### Bottom-Up Validation

Pairs of protein accession numbers were submitted to PPI-ID. A prediction was considered successful if the proper interaction interface was identified in PPI-ID output. PPI-ID bottom-up validation yielded respective accuracy rates of 98% and 95% for DDIs and DMIs. The missing DDI and DMI identifications arose because of missing Pfam or SLiM identifiers in the InterPro and ELM databases.

### Using PPI-ID with PAE Viewer to enhance understanding of motifs in IDRs

Another AlphaFold product that can be evaluated to ascertain the quality of a PPI model is Predicted Aligned Error (PAE). PAE expresses the error in angstroms of the relative position of two amino acids. A low error within interdomain quadrants of PAE plots, which are quadrants 1 and 3 (top right and bottom left quadrants; Figure 4), suggests a physical interaction between the proteins (18). The PAE Viewer web tool plots PAE values and lets one determine the residue numbers of all plotted points, a feature that complements PPI-ID (https://subtiwiki.uni-goettingen.de/v4/paeViewerDemo) (19). If a PPI-ID predicted DDI or DMI were valid, one would expect the predicted interface to have a very low PAE score. In turn, one can determine whether regions of low PAE value house known interaction domains or motifs. Such outcomes would be supportive that the interaction is real.

**Figure 4.**
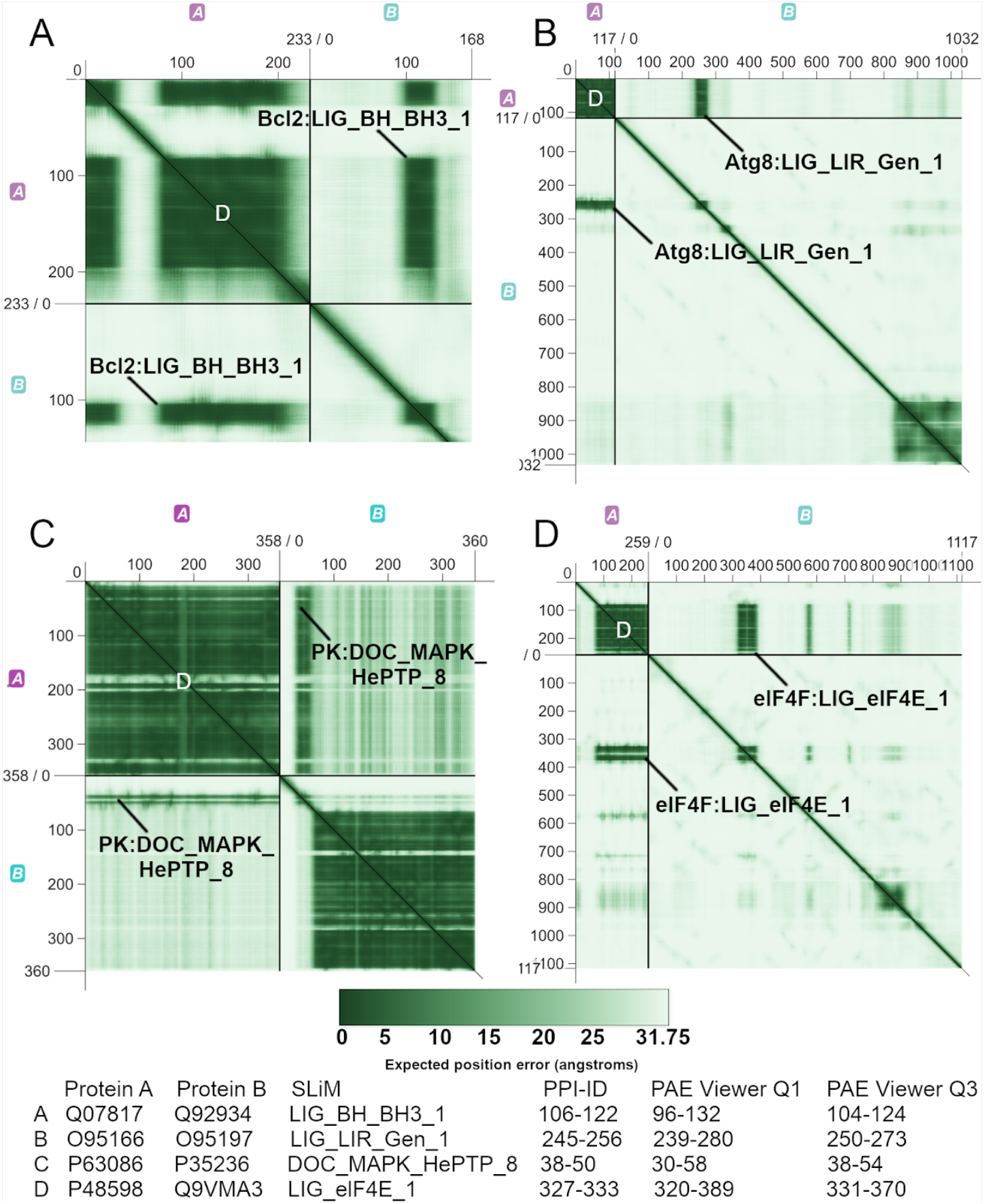
PPI-ID in conjunction with PAE Viewer enhances confidence in the protein interactions visualized in a model. Panels are PAE plots of DMI protein pairs. Within each plot, quadrant 1 (Q1) is the top right hand quadrant and quadrant 3 (Q3) is the bottom left quadrant. Q1 and Q3 represent interactions between a motif in protein B and a domain in protein A. The heat map represents the positional error between amino acid pairs measured in angstroms. PAE Viewer reports residue numbers from positions on the plot. Using these, we correlated the outputs of PPI-ID and PAE Viewer. The PPI-ID contact distance filter was set to ≤ 5 angstroms. SLiMs identified by PPI-ID are labeled. The predicted domain that the SLiM interacts with is the dark green square to the left of the labeled Q1 SLiM and above the labeled Q3 SLiM. The domains, identified by the white D in panels A-D, are the Bcl-2 apoptosis regulator domain (Bcl-2), autophagy protein Atg8 ubiquitin like (Atg8), protein kinase domain (PK), and eukaryotic initiation factor 4E domains (eIF4E), respectively. The table shows residue numbers of SLiMs identified by PPI-ID and the PAE Viewer identified residues with low PAE scores (dark stripes) in Q1 and Q3. The programs show good agreement in identification of the same sequences based on distinct criteria.

The protein pairs in Figure 4 were processed with PPI-ID, which mapped a SLiM inside an IDR of protein B and predicted that the SLiM is within 5 angstroms of the target domain within protein A (labeled in Figure 4). PPI-ID mapped interacting protein regions solely based on information about, and the distances between, domains and motifs. PPI-ID provided the residue numbers for the DMIs, and PAE Viewer provided residue numbers of the regions of low PAE within the IDR of protein B. We used these residue numbers to correlate the information from the PAE plot to the output from PPI-ID. In Figure 4, a DMI is predicted to occur at the sites of low relative positional error between the proteins, bolstering the evidence that PPI-ID provides a salient interpretation of the protein motifs documented within the SLiM database provided by ELM.

## DISCUSSION

AlphaFold-Multimer is an important tool for advancing our understanding of how proteins interact (5, 6). PPI-ID is a program that can be run from its website or downloaded and run locally. It identifies potential interaction interfaces between proteins and characterizes interaction interfaces in AlphaFold-produced models. The top-down approach (identification of potential DDIs and DMIs that are in close proximity) performed less well than the bottom-up approach (simple search of protein sequences for compatible DDI and DMI pairs). The reduced top-down accuracy is attributable to AlphaFold-Multimer error when folding larger proteins or protein complexes, as larger proteins are more susceptible to interface misidentification (Supplemental data). Both Bret et al. (7) and Lee et al. (8) have observed similar declines in AlphaFold-Multimer accuracy as protein sizes increase. Bret et al. (7) suggest that diminished accuracy in larger complexes arises from intramolecular contacts that mask correct domain/motif interaction. Additionally, top-down performance worsens when attempting to predict PPIs between two polypeptides that are part of a complex composed of more than two proteins. The four instances in which a DDI was not correctly predicted in the Top Down Approach involved proteins that are part of a complex of three or more proteins. This may result from the fact that multimeric protein complexes often are synergistically held together by each constituent polypeptide chain, and missing polypeptides may lead to the absence of PPIs essential for proper complex assembly.

PPI-ID is a convenient tool for scanning a pair of proteins or a model of a protein dimer for protein domains and motifs that are annotated as interacting in well-curated standard databases. The veracity of PPI-ID mapping of DDIs and DMIs is dependent on the quality of the domain and motif databases and the quality of the protein model. Since domain and motif databases and protein modeling fidelity continuously improve, we expect PPI-ID performance to also improve over time. We recognize that for any given protein-protein interaction prediction, it would be best to confirm the interaction by a physical technique. However, interactive rapid screening, as that provided by PPI-ID, can help focus one on the most likely interaction candidates.

PPI-ID was developed to screen through a list of PPI candidates and is particularly useful when used in conjunction with PAE Viewer (19). Using both programs in tandem enables the correlation of interaction domains and motifs with regions of low positional error. Such analysis provides additional support for the idea that the identified domains and motifs of interest stabilize one another and drive a protein:protein association.

Another program focused on interpreting SLiMs is SLiMAn (20). The unique capabilities of SLiMAn include interactome-level analysis of DMIs and the filtering for specific properties, e.g., within disordered regions or post-translational modifications. On the other hand, the unique capabilities of PPI-ID include the prediction of both DDIs and DMIs, PPI visualization that is granular and interactive, and filtering protein:protein interfaces based on their intermolecular distance. SLiMAn seems well suited for characterizing a known protein interactome while PPI-ID is best suited for providing evidence of whether protein pairs belong to the same interactome. We imagine that sequential use of SLiMAn and PPI-ID could have utility.

PPI-ID is capable of visualizing protein:protein interfaces, filtering interfaces by contact distance, and residue labeling. Furthermore, PPI-ID performs DDI and DMI prediction from protein sequence information. PPI-ID is a compact and flexible tool that streamlines the processes of both PPI structural analysis and protein sequence delimitation for structural prediction.

## DATA AVAILABILITY

All program code and data have been archived under the FigShare DOI of 10.6084/m9.figshare.28266584.

## SUPPLEMENTARY DATA

Supplementary Table 1 and Supplementary Table 2

## AUTHOR CONTRIBUTIONS

Haley Goodwin: Conceptualization, Data curation, Methodology, Software, Formal analysis, Validation, Funding acquisition, Writing—original draft, review & editing. Nigel Atkinson: Conceptualization, Validation, Funding acquisition, Supervision, Writing—original draft, review & editing, Resources, Project administration.

## ACKNOWLEDGEMENTS

The authors acknowledge the Texas Advanced Computing Center (TACC) at The University of Texas at Austin for providing computational resources that have contributed to the research results reported within this paper. URL: http://www.tacc.utexas.edu. The authors also thank Mr. James R. Derry for his assistance helping us navigate the universities web services.

